# Metagenomic Nanopore sequencing of influenza virus direct from clinical respiratory samples

**DOI:** 10.1101/676155

**Authors:** Kuiama Lewandowski, Yifei Xu, Steven T. Pullan, Sheila F. Lumley, Dona Foster, Nicholas Sanderson, Alison Vaughan, Marcus Morgan, Nicole Bright, James Kavanagh, Richard Vipond, Miles Carroll, Anthony C. Marriott, Karen E Gooch, Monique Andersson, Katie Jeffery, Timothy EA Peto, Derrick W. Crook, A Sarah Walker, Philippa C. Matthews

## Abstract

Influenza is a major global public health threat as a result of its highly pathogenic variants, large zoonotic reservoir, and pandemic potential. Metagenomic viral sequencing offers the potential of a diagnostic test for influenza which also provides insights on transmission, evolution and drug resistance, and simultaneously detects other viruses. We therefore set out to apply Oxford Nanopore Technology to metagenomic sequencing of respiratory samples. We generated influenza reads down to a limit of detection of 10^2^-10^3^ genome copies/ml in pooled samples, observing a strong relationship between the viral titre and the proportion of influenza reads (p = 4.7×10^-5^). Applying our methods to clinical throat swabs, we generated influenza reads for 27/27 samples with high-to-mid viral titres (Cycle threshold (Ct) values <30) and 6/13 samples with low viral titres (Ct values 30-40). No false positive reads were generated from 10 influenza-negative samples. Thus Nanopore sequencing operated with 83% sensitivity (95% CI 67-93%) and 100% specificity (95% CI 69-100%) compared to the current diagnostic standard. Coverage of full length virus was dependent on sample composition, being negatively influenced by increased host and bacterial reads. However, at high influenza titres, we were able to reconstruct >99% complete sequence for all eight gene segments. We also detected Human Coronavirus and generated a near complete Human Metapneumovirus genome from clinical samples. While further optimisation is required to improve sensitivity, this approach shows promise for the Nanopore platform to be used in the diagnosis and genetic analysis of influenza and other respiratory viruses.

## Introduction

Influenza A is an RNA Orthomyxovirus of approximately 13Kb length, with an eight-segment genome. It is typically classified on the basis of Haemagglutinin (HA) and Neuraminidase (NA), of which there are 16 and 9 main variants respectively [1]. Genetic reassortment underpins the potential for transmission between different host species [2], and for the evolution of highly pathogenic variants [3–6], recognised in the WHO list of ‘ten threats to global health’ [7]. Seasonal influenza causes an estimated 650,000 deaths globally each year, and H3N2 alone kills 35,000 each year in the USA [1,8]. Certain groups are particularly at risk, including older adults, infants and young children, pregnant women, those with underlying lung disease, and the immunocompromised [9]. The burden of disease disproportionately affects low/middle income settings [10]. Influenza diagnostics and surveillance are fundamental to identify the emergence of novel strains, to improve prediction of potential epidemics and pandemics [4,11], and to inform vaccine strategy [12]. Diagnostic data facilitate real-time surveillance, can underpin infection control interventions [13,14] and inform the prescription of Neuraminidase Inhibitors (NAI) [9].

Currently, most clinical diagnostic tests for influenza depend on detecting viral antigen, or PCR amplification of viral nucleic acid derived from respiratory samples [15]. These two approaches offer trade-offs in benefits: antigen tests (including point-of-care tests (POCT)) are typically rapid but low sensitivity [16–18], while PCR is more time-consuming but more sensitive [9]. Irrespective of test used, most clinical diagnostic facilities report a non-quantitative (binary) diagnostic result, and the data generated for influenza have limited capacity to inform insights into epidemiological linkage, vaccine efficacy or anti-viral susceptibility. Given this, there is an aspiration to generate new diagnostic tests that combine speed (incorporating potential for POCT [19,20]), sensitivity, detection of co-infection [21,22], and generation of quantitative or semi-quantitative data that can be used to identify drug resistance and reconstruct phylogeny to inform surveillance, public health strategy, and vaccine design.

The application of Oxford Nanopore Technologies (ONT; https://nanoporetech.com/) to generate full-length influenza sequences from clinical respiratory samples can address these challenges. ONT offers a ‘third-generation’, portable, real-time approach to generating long-read sequence data, with demonstrated success across a range of viruses [21,23–25]. To date, Nanopore sequencing of influenza has been reported using high titre virus from an *in vitro* culture system, producing full length genome sequences through direct RNA sequencing [26] or targeted enrichment by either hybridisation of cDNA [27] or influenza-specific PCR amplification [28].

We therefore aimed to optimise a metagenomic protocol for detecting influenza viruses directly from clinical samples using Nanopore sequencing. We determine its sensitivity compared to existing diagnostic methods and its accuracy compared to short-read (Illumina) sequencing, using clinical samples from hospital patients during an influenza season, and samples from a laboratory controlled infection in ferrets.

## Results

### Method optimisation to increase proportion of viral reads derived from throat swabs

We first sequenced five influenza A positive and five influenza-negative throat swabs, each spiked with Hazara virus control at 10^4^ genome copies/ml. Using the SISPA approach [21] followed by Nanopore sequencing, we produced metagenomic data dominated by reads that were bacterial in origin, with extremely few viral reads detected. Passing the sample through a 0.4 μm filter prior to nucleic acid extraction increased the detection of viral reads by several orders of magnitude (Fig S1). Filtration is relatively expensive, so we also assessed the alternative approach of adding a rapid centrifugation step to pellet bacterial and human cells, followed by nucleic acid extraction from the supernatant. We used a pooled set of influenza A positive samples (concentration 10^6^ genome copies/ml), to provide a large enough sample to assess reproducibility, with the Hazara control spiked in at 10^4^ genome copies/ml. Enrichment for influenza and Hazara was similar for filtration vs centrifugation, based on reads mapping to the viral genome (Fig S2). As centrifugation is simpler and cheaper, we selected this approach for all further testing.

### Method optimisation to reduce time for cDNA synthesis

Synthesis of tagged, randomly primed cDNA and its subsequent amplification via SISPA [21] required lengthy reverse transcription and PCR steps (1h and 3h 45min) respectively. Optimising these stages upgraded the reverse transcriptase from SuperScriptIII to SuperScriptIV (ThermoFisher), reduced incubation time to 10min (processing time reduction of 50min) and reduced the PCR extension time within each cycle from 5min to 2min (1h 30min processing time reduction). Comparing this final method with our original protocol, using triplicate extractions from the pooled set of influenza A positive samples, demonstrated no significant loss in performance in the more rapid protocol (Fig S3) and we adopted this approach as our routine protocol.

### Consistent retrieval of Hazara virus by Nanopore sequencing

Starting with an influenza A positive sample pool (10^6^ genome copies/ml), we made three volumetric dilution series using three independent influenza-negative pools (Fig 1). The total quantity of cDNA after preparation for sequencing was consistently higher in all samples using negative pool-3 as diluent (Fig 2A), indicating the presence of a higher concentration of non-viral RNA within pool-3. This is likely due to host cell lysis or higher bacterial presence, and demonstrates the variable nature of throat swab samples.

**Figure 1:**
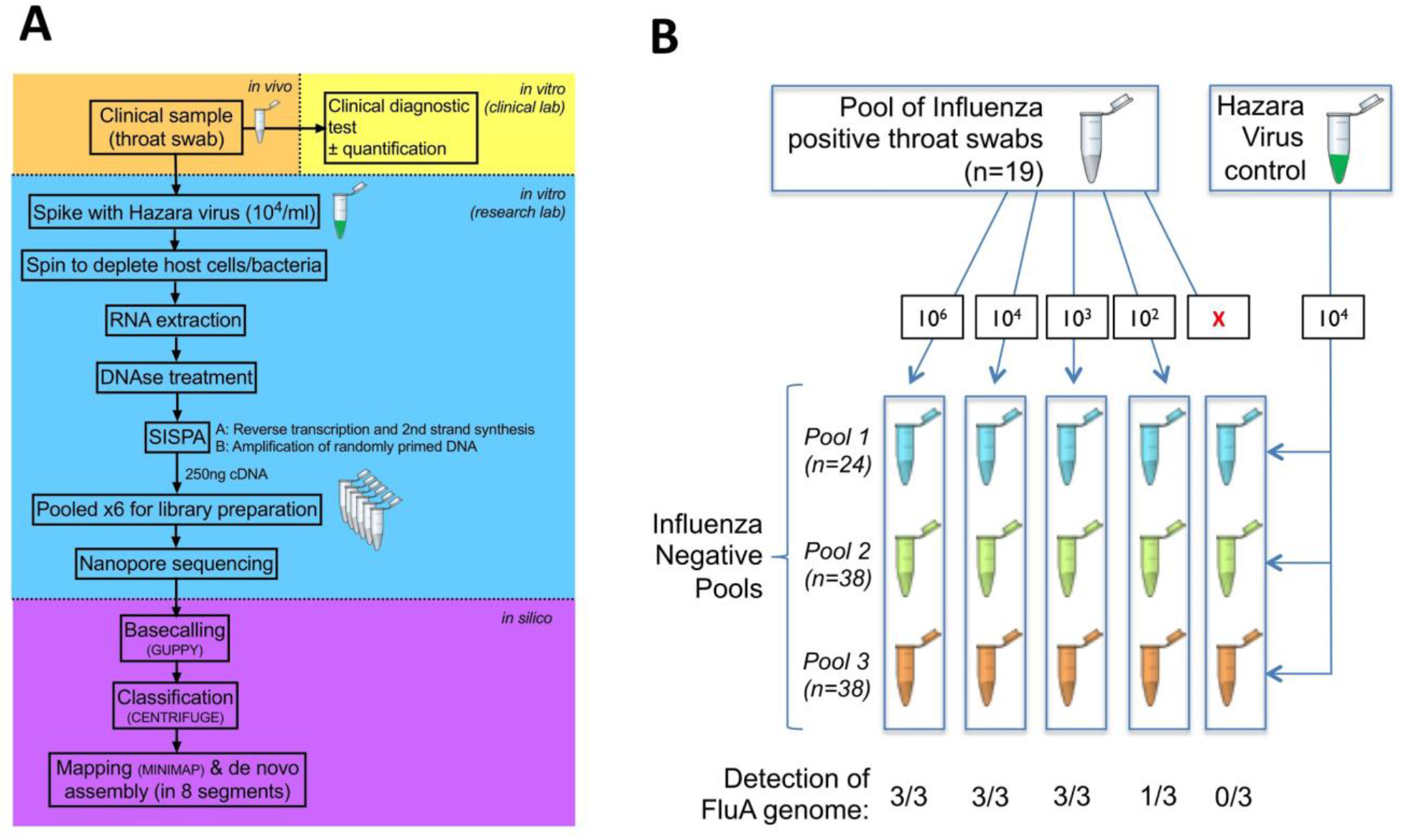
Schematic to show processing protocol through clinical and research pipelines for influenza diagnosis. (A) Clinical sample collection (orange), clinical diagnostic testing (yellow), sample processing and sequencing using Oxford Nanopore Technologies (blue), processing of sequence data (purple). (B) Outline of pooled influenza-positive samples into influenza-negative background, to generate varying titres of influenza virus (from zero up to 10^6^ genome copies/ml), undertaken in triplicate, and spiked with a standard titre of Hazara virus control at 10^4^ genome copies/ml).

**Figure 2.**
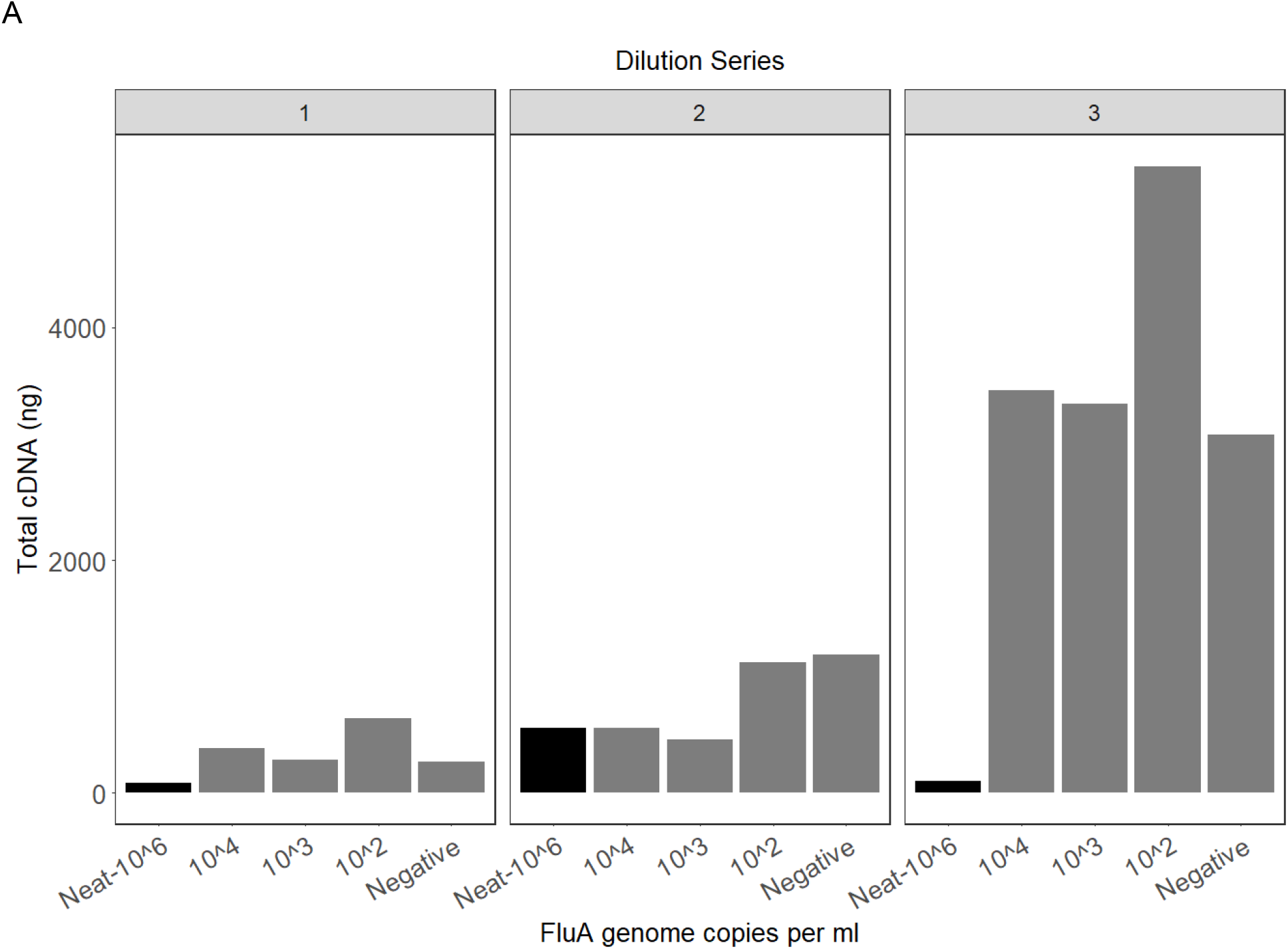
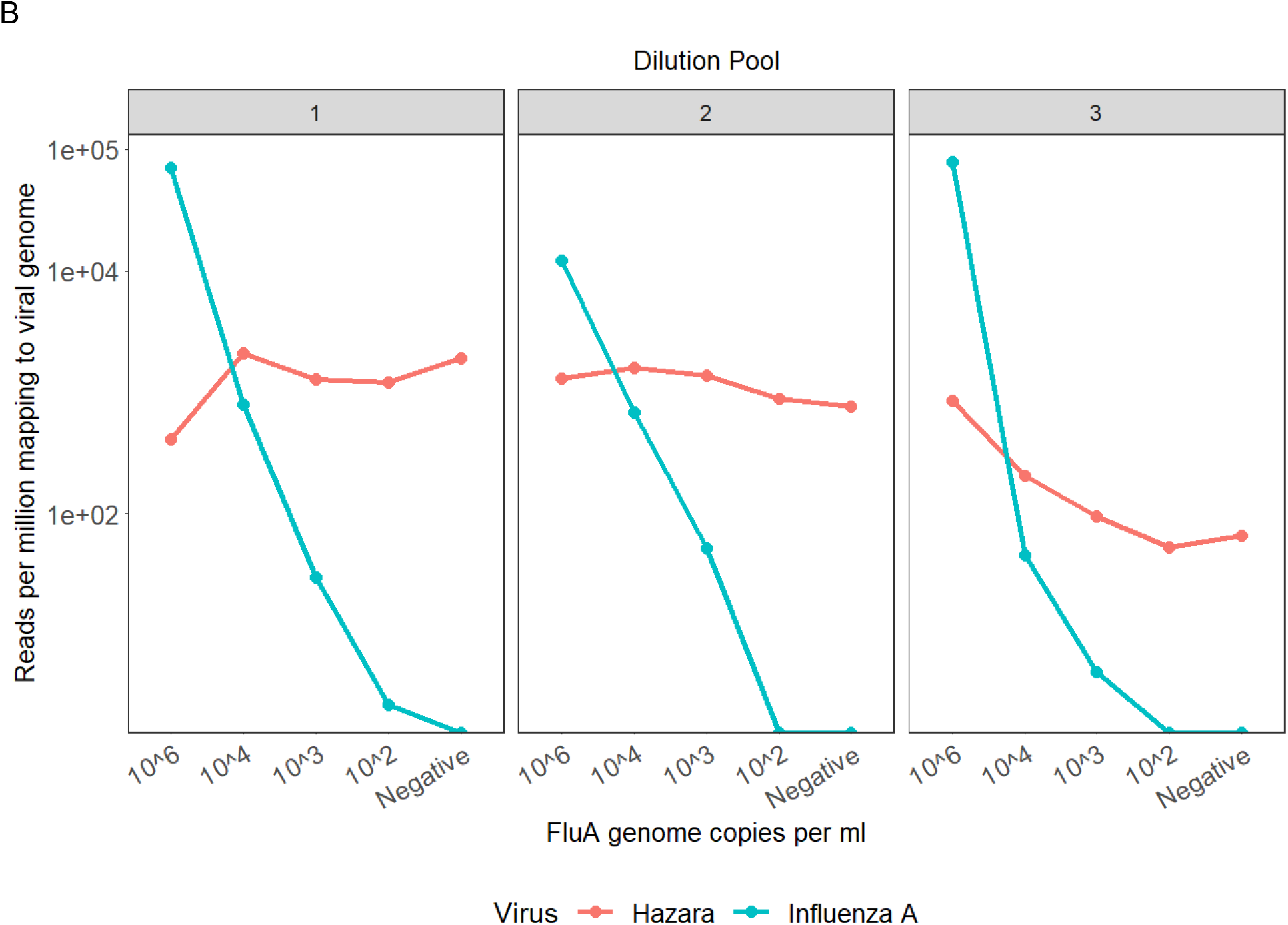
Characteristics of three pools of influenza-negative throat swabs, and Nanopore sequence results following spiking with influenza A. (A) Total concentration of cDNA produced per pooled sample following amplification by the SISPA reaction, grouped by dilution series. The 10^6^ sample in each pool is the original, undiluted material, represented by the black filled bars. Samples diluted to influenza titres of 10^4^, 10^3^ and 10^2^ contain more cDNA due to higher background material (bacterial/human) present in the diluent. Dilution series 1 and 2 contain comparable amounts of background material; dilution series 3 contains substantially more background. (B) Viral reads generated by Nanopore sequencing of samples with different titres of influenza A, and a consistent titre of Hazara virus (10^4^ genome copies/ml). Graphs show reads per million of total reads mapping to influenza A or Hazara virus genomes, across the three individual dilution series. Note logarithmic scale on y-axis.

We consistently retrieved Hazara virus reads from all three dilution series by Nanopore sequencing, independently of influenza virus titre in the sample (Fig 2B). Sequencing from dilution series 1 and 2 gave a consistent proportion of total reads mapping to the Hazara genome, across dilutions and between the first two pools with mean values per pool of 1.4×10^3^ ±660 RPM (reads per million (of total reads) ± standard deviation) and 1.2×10^3^ ±350 RPM respectively. The pool-3 dilution series generated 260 ±340 RPM Hazara reads across samples, and showed a decreasing trend associated with increased dilution factor, as increasingly more non-viral RNA was introduced from this high-background pool.

### Limit of influenza virus detection by Nanopore from pooled samples

Nanopore sequencing of the triplicate SISPA preparations of the influenza A positive pool produced mean 5.3×10^4^ ±3.6×10^4^ RPM mapping to the influenza A genome (Fig 2B). Across the dilution series, the proportion of influenza reads was strongly associated with influenza virus titre (p-value = 4.7×10^-5^), but was also influenced by which negative pool was used for dilution, consistent with the pattern observed for the Hazara control. Sequencing the negative controls (pools with no influenza virus spike) generated no reads mapping to influenza. At influenza virus titres <10^3^ copies/ml, influenza reads were inconsistently detected across the samples (Fig 2B), suggesting the limit of detection is between 10^2^-10^3^ influenza copies/ml.

### Retrieval and reconstruction of complete influenza genomes from pooled/spiked samples

For the Hazara virus control (10^4^ genome copies/ml spike), genome coverage was 81.4-99.4% (at 1x depth) for pools 1 and 2. Coverage in the high-background pool 3 was more variable (21.5-96.5%; Fig 3A). Influenza A genome coverage at 10^6^ copies/ml was ≥99.3% for each segment in all samples (Fig 3A). At 10^4^ genome copies/ml of influenza, mean 1x coverage per segment was 90.3% for pools 1 and 2, but was substantially reduced in the high-background pool 3 to 5.7% (Fig 3A). At influenza titres <10^4^ copies/ ml, coverage was highly variable across genome segments. However, when present at 10^3^ copies/ml, 2/3 pools had sufficient data for correct subtyping as H3N2 (Table 1).

**Table 1.**
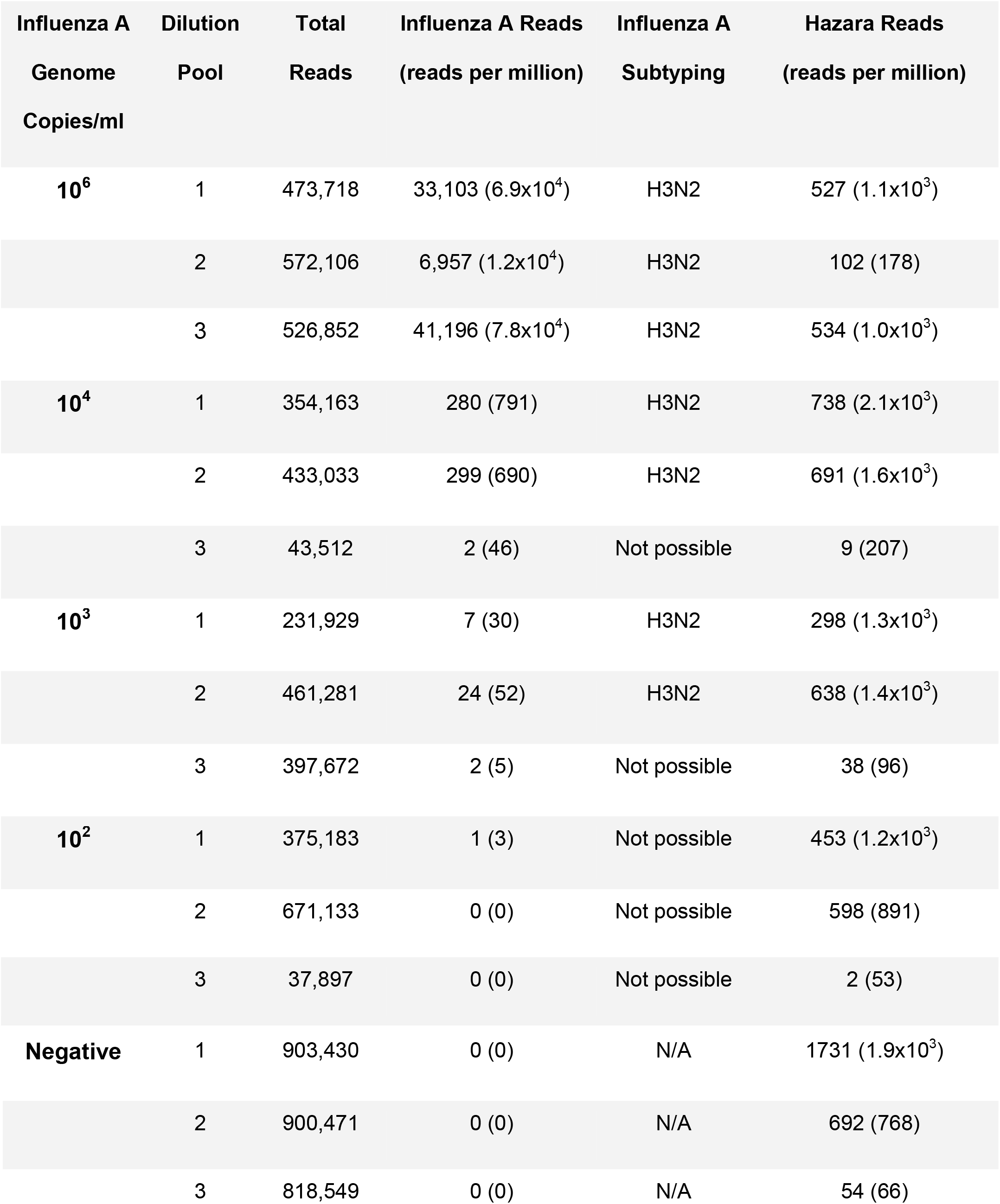
Summary of results from Nanopore sequencing based on pooled samples with varying titres of Influenza A, and a consistent titre of Hazara virus control. Each dilution is undertaken in triplicate (shown as 3 dilution pools).

| **Influenza A**<br>**Genome**<br>**Copies/ml** | **Dilution**<br>**Pool** | **Total**<br>**Reads** | **Influenza A Reads**<br>**(reads per million)** | **Influenza A**<br>**Subtyping** | **Hazara Reads**<br>**(reads per million)** |
| --- | --- | --- | --- | --- | --- |
| **10<sup>6</sup>** | 1 | 473,718 | 33,103 (6.9x10<sup>4</sup>) | H3N2 | 527 (1.1x10<sup>3</sup>) |
|  | 2 | 572,106 | 6,957 (1.2x10<sup>4</sup>) | H3N2 | 102 (178) |
|  | 3 | 526,852 | 41,196 (7.8x10<sup>4</sup>) | H3N2 | 534 (1.0x10<sup>3</sup>) |
| **10<sup>4</sup>** | 1 | 354,163 | 280 (791) | H3N2 | 738 (2.1x10<sup>3</sup>) |
|  | 2 | 433,033 | 299 (690) | H3N2 | 691 (1.6x10<sup>3</sup>) |
|  | 3 | 43,512 | 2 (46) | Not possible | 9 (207) |
| **10<sup>3</sup>** | 1 | 231,929 | 7 (30) | H3N2 | 298 (1.3x10<sup>3</sup>) |
|  | 2 | 461,281 | 24 (52) | H3N2 | 638 (1.4x10<sup>3</sup>) |
|  | 3 | 397,672 | 2 (5) | Not possible | 38 (96) |
| **10<sup>2</sup>** | 1 | 375,183 | 1 (3) | Not possible | 453 (1.2x10<sup>3</sup>) |
|  | 2 | 671,133 | 0 (0) | Not possible | 598 (891) |
|  | 3 | 37,897 | 0 (0) | Not possible | 2 (53) |
| **Negative** | 1 | 903,430 | 0 (0) | N/A | 1731 (1.9x10<sup>3</sup>) |
|  | 2 | 900,471 | 0 (0) | N/A | 692 (768) |
|  | 3 | 818,549 | 0 (0) | N/A | 54 (66) |

**Figure 3.**
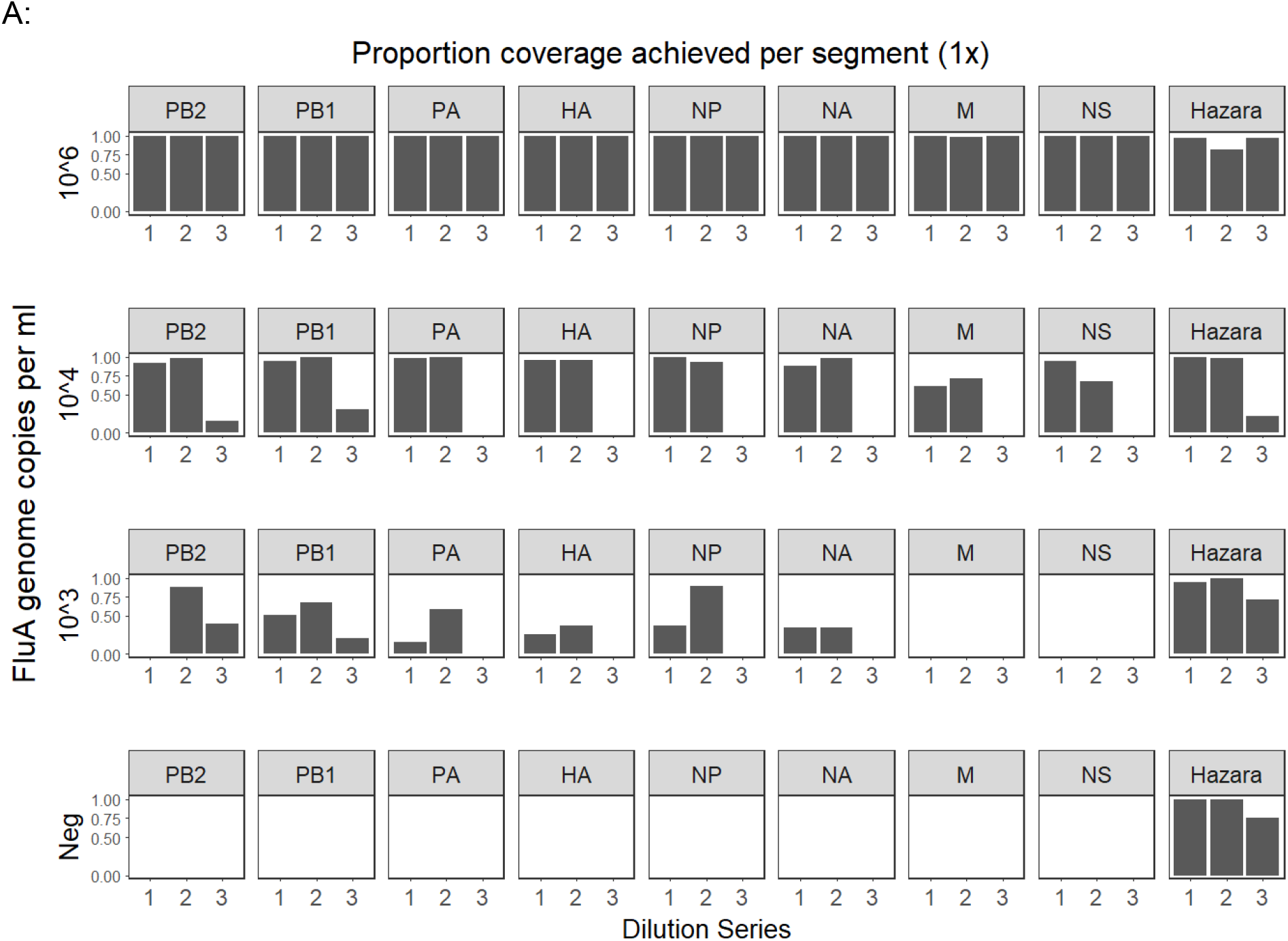
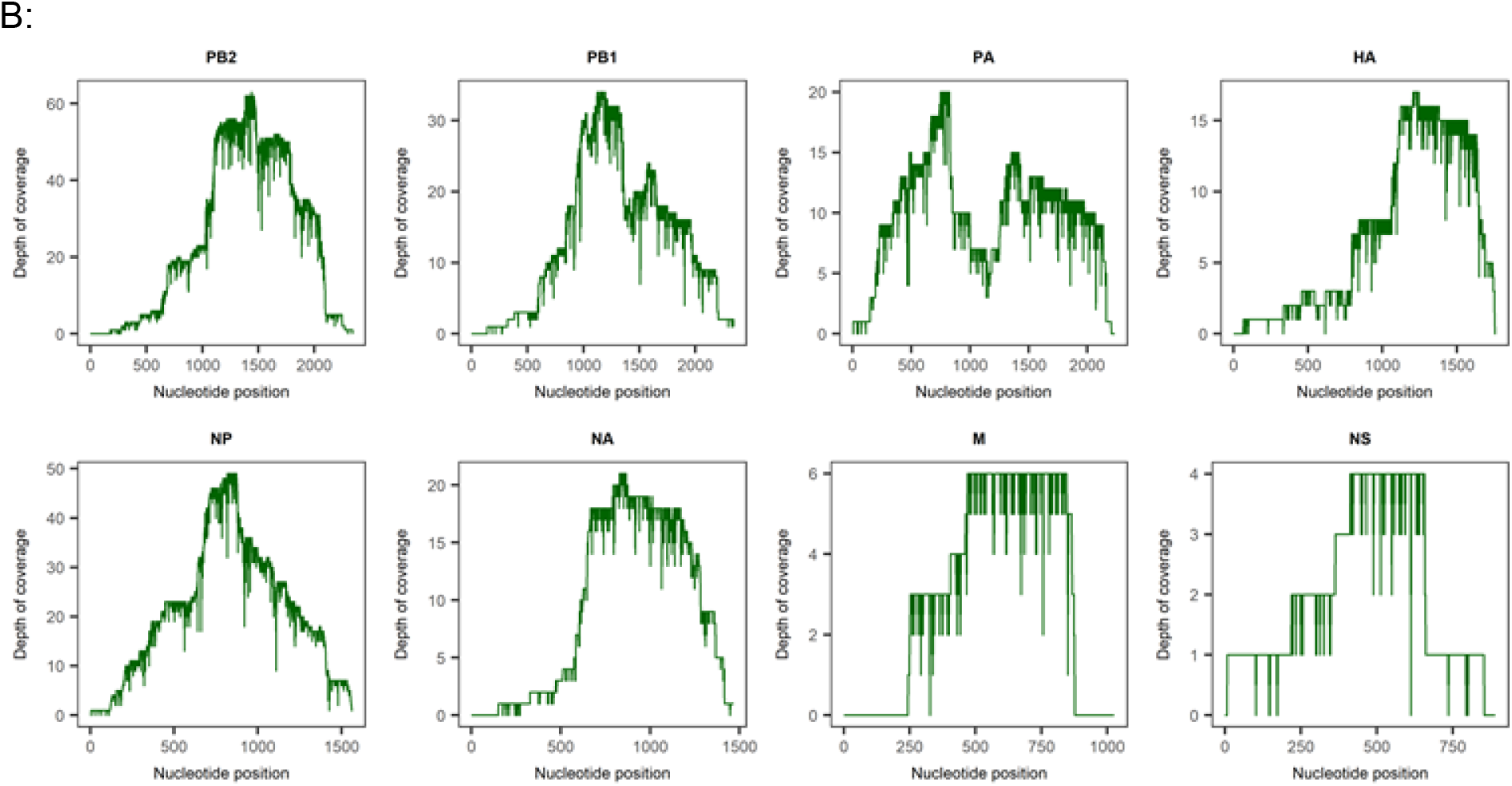
Coverage of influenza and Hazara genome segments achieved by Nanopore sequencing from pooled samples. (A) Data from three dilution series of pooled influenza positive samples, diluted with three separate negative sample pools to generate different titres of influenza virus. Each individual dilution was spiked with Hazara at 10^4^ genome copies/ml. Proportion of genome covered at 1x depth is shown for each of the eight influenza genome segments (encoding: PB2 - polymerase subunit, PB1 - polymerase subunit, PA - polymerase acidic protein, HA - hemagglutinin, NP- nucleocapsid protein, NA - neuraminidase, M - matrix proteins, NS - nonstructural proteins) across the three dilution series. Coverage of the Hazara genome is plotted as total of all three genome segments for simplicity. (B) Representative coverage plot of influenza A genome segments from the dilution series 1 sample at 10^4^ influenza copies per ml.

### Influenza detection from individual clinical samples

Having demonstrated our ability to retrieve influenza sequences from pooled influenza-positive material diluted with negative samples, we next applied our methods to individual anonymised clinical samples, 40 testing influenza-positive and 10 influenza-negative in the clinical diagnostic laboratory. Data yield varied between flow cells (range 2.5×10^6^ - 13.2×10^6^ reads from up to 12 multiplexed samples). Within flow cells, barcode performance was inconsistent when using a stringent, dual barcode, demultiplexing method [22]. From each clinical sample, the range of total reads generated was 1.0×10^5^ to 2.4×10^6^ (median 3.8×10^5^) (Table S1).

Reads mapping to either influenza A or B genomes were present in all 27 samples with Cycle threshold (Ct) <30 (range 6 to 274,955 reads). At Ct >30, 6/13 samples generated influenza reads (range 6 to 92,057), (difference between sensitivity at a Ct threshold of 30, p<0.0001, Fig 4) The highest Ct value at which any influenza reads were detected was Ct 36.8 (Sample 37; 17 reads influenza A). No reads classified as influenza were obtained from sequencing the ten GeneXpert-negative samples (Table S1). Based on this small dataset, sensitivity is 83% and specificity 100% (95% CI 67-93% and 69-100%, respectively). There was a strong correlation between Ct value and both reads per sample classified as influenza (R-squared =0.604) and number of influenza reads per million reads (R-squared =0.623) (Fig 4).

**Figure 4:**
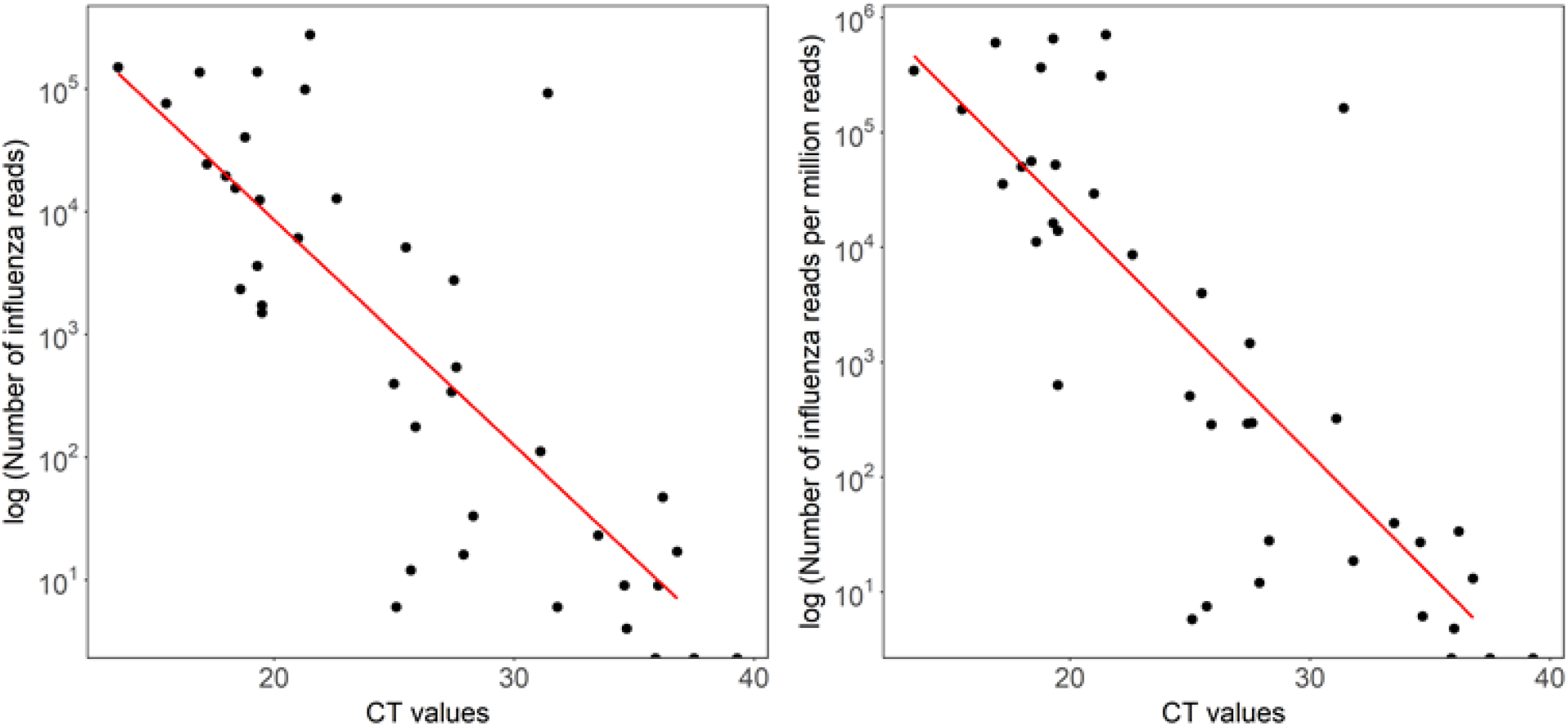
Total and proportion of influenza reads derived by Nanopore sequencing of individual samples across a range of Ct values. Ct values were derived by testing using GeneXpert (Cepheid) in a clinical diagnostic laboratory. (A) Correlation between Ct value and total number of influenza reads generated. R^2^= 0.604, p-value = 2.47e-08. (B) Correlation between Ct value and number of influenza reads per million reads. R^2^= 0.623, p-value = 1.07e-08.

### Detection of Hazara virus internal control

Detection of the control virus (Hazara at 10^4^ genome copies/ml) was highly variable, demonstrating that levels of background non-target RNA are a major source of inter-sample variation. Hazara reads per sample ranged from 0 to 13.5×10^3^ (0-3.5×10^4^ RPM) with a median of 70 (160 RPM) and mean of 706 (1.7×10^3^ RPM) (Table S1). Four (8%) of 50 samples generated no detectable Hazara reads, two with high numbers of influenza reads (Sample 6: Ct 18.4, 1.5×10^4^ influenza A reads and Sample 1: Ct 13.5, 1.5×10^5^ influenza B reads) acting to dilute the control signal. The other two samples contained no detectable influenza reads (Sample 34: Ct 35.9, Sample 46, Flu negative). The lack of control detection therefore indicates a loss of assay sensitivity due to high levels of background nucleic acid present in some samples.

### Comparison with Illumina sequencing

A subset of 15 samples were selected from across the viral titre range and re-sequenced on an Illumina MiSeq. Proportions of reads generated that mapped to the influenza genome were similar across the two sequencing technologies (Figure S4).

### Influenza phylogeny and typing

We reconstructed the phylogeny using consensus sequences for the HA gene (Fig 5). This demonstrates closely related sequences, as expected within one location in a single influenza season, and complete concordance between the sequences obtained by Nanopore vs Illumina for the subset of isolates sequenced by both technologies.

**Figure 5:**
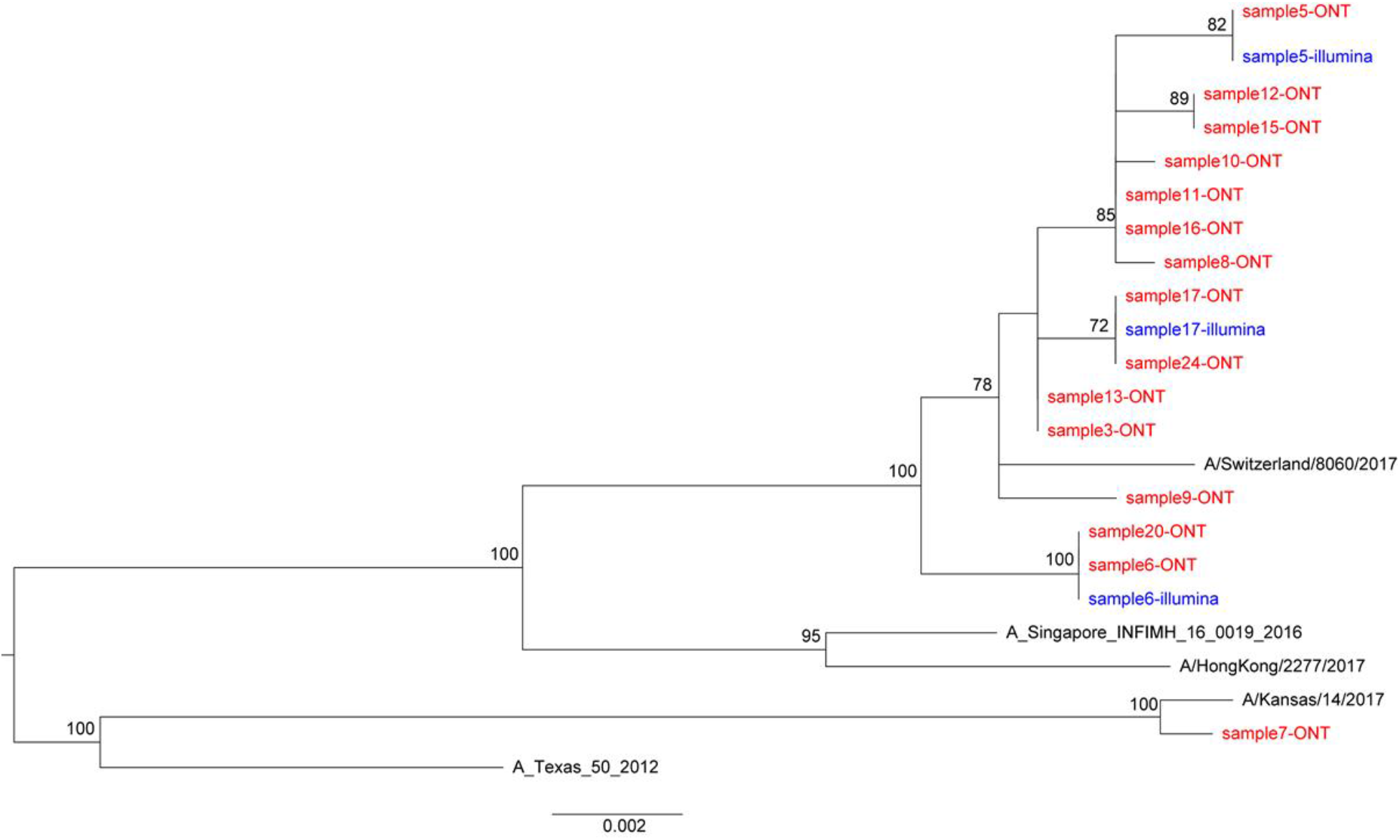
Phylogenetic trees of consensus Influenza HA gene derived by Nanopore and Illumina sequencing. Maximum likelihood tree generated using 500 bootstrap replicates in RAxML v8.2.10 software. Bootstrap values >70 are shown. Scale bar shows substitutions per site. Red and blue indicate sequences derived from Nanopore and illumina sequencing, respectively. References sequences are shown in black.

### Detection of other RNA viruses in clinical samples

Within the 50 clinical samples sequenced, we found limited evidence of other RNA viruses. Sample 6 produced 109 reads mapping to Human Coronavirus in addition to >1.5×10^4^ influenza A reads, suggesting co-infection. We also derived >4.0×10^4^ reads from Human Metapneumovirus from an influenza-negative sample, providing a near complete (99.8%) genome from one sample (Figure S1, Sample I), further detailed here [29].

### Animal time-course study

Finally, we used samples collected from a previous animal experiment [30] to test the reproducibility of our methods across a time-course model of influenza A infection (three ferrets pre-infection (day -3) and then at days 1, 2, 3 and 5 following laboratory infection with influenza A). The proportion of viral reads present at each timepoint was highly congruent with viral titre (titre shown in Fig 6A and sequencing reads in Fig 6B). Consensus genome sequences were generated from days two, three and five and were 100% concordant with Illumina-derived consensus sequences from the same cDNA.

**Figure 6:**
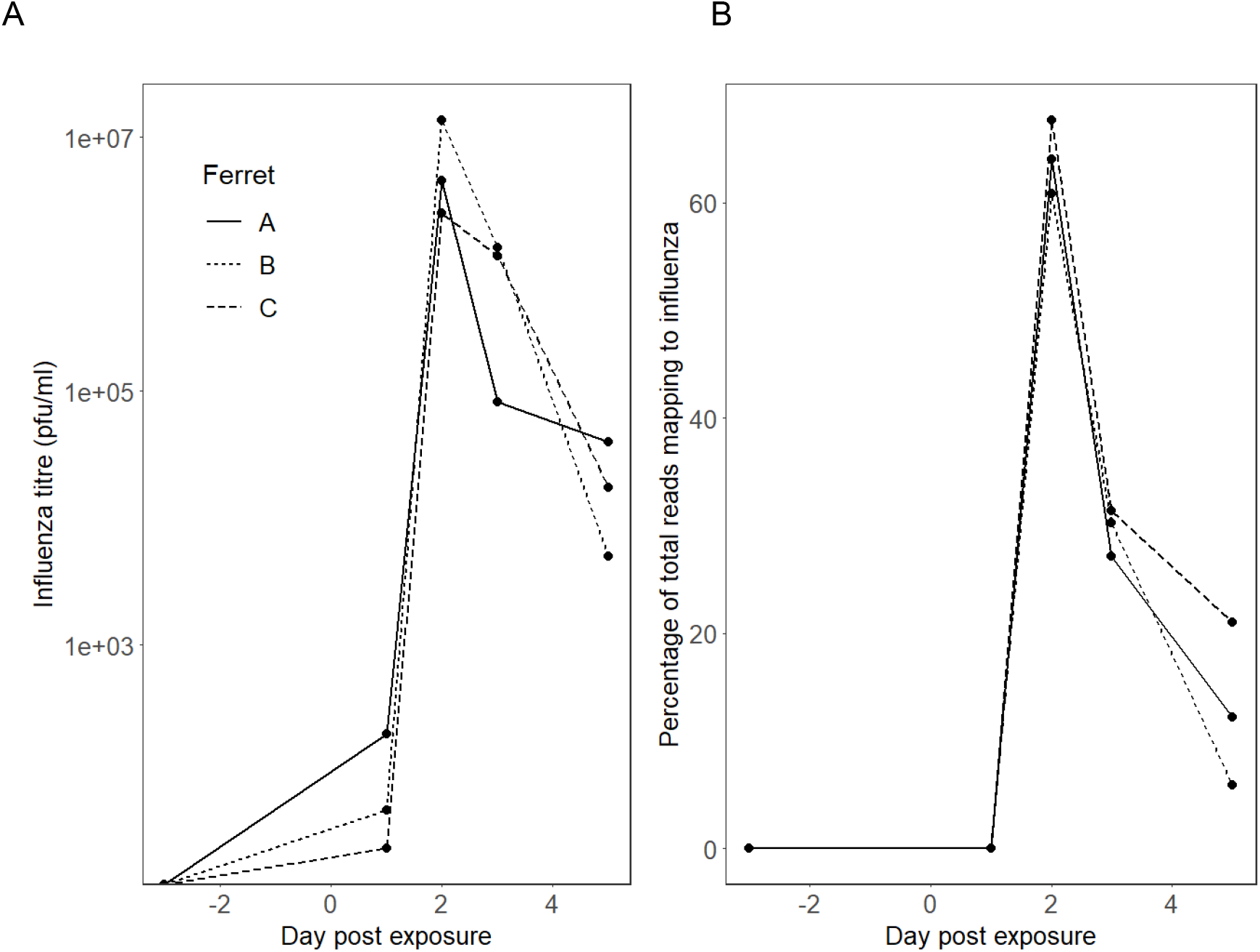
Time course experiment showing influenza A infection in three laboratory ferrets. Infection was introduced at day 0. Samples were collected three days prior to infection, and at days 1, 3 and 5 post infection. (A) Influenza titre (log scale) and (B) proportion of total nanopore reads (linear scale) mapping to influenza A from metagenomic sequencing of ferret nasal washes taken pre and post influenza challenge.

## DISCUSSION

To our knowledge, this is the first report of successfully applying metagenomic Nanopore sequencing directly to respiratory samples to detect influenza virus and generate influenza sequences. The approach demonstrates excellent specificity; sensitivity varies by viral titre, but is comparable to existing laboratory diagnostic tests for Ct values <30. Our optimised protocol depletes human and bacterial nucleic acids, and reduces the time from sample to sequence. This method has the potential to be further optimised and validated to improve sensitivity for influenza, identify other RNA viruses, detect drug-resistance mutations, and provide insights into quasispecies diversity [31,32]. At a population level, this sequence-based diagnostic data can, in addition, provide phylo-epidemiological reconstruction, insights into transmission events, potential for estimating vaccine efficacy [33], and approaches for public health intervention [34].

Whole genome viral sequencing can accurately trace outbreaks across time and space [35]. The metagenomic method employed here produced >90% complete genomes for 17/27 samples with a Ct value ≤30, demonstrating the ability of metagenomics to produce sufficient data for influenza diagnostics and genome characterisation, whilst also detecting and sequencing other common RNA viruses.

Despite time reductions in wet laboratory processing, this method requires further modification to simplify and accelerate the protocol if it is to become viable as a near-to-patient test. High error rates are a recognised concern in Nanopore sequence data, and cross-barcode contamination can create challenges when low and high titre samples are batched together [22]. To avoid these problems, we batched samples according to Ct value, and applied stringent barcode demultiplexing criteria; however, this reduces the total data available for analysis [22]. For future primary diagnostic use it would be preferable to sequence samples individually using a lower throughput flow cell e.g. ONT Flongle. Careful optimisation of laboratory and bioinformatic methods are required to resolve individual sequence polymorphisms, particularly for drug resistance alleles.

IDSA guidelines [9] recommend nasal/nasopharyngeal specimens for influenza diagnosis, but throat swabs are easier to collect in clinical practice and therefore accounted for the majority of our diagnostic samples. Further work is needed to investigate the sensitivity and specificity of our protocol for a wider array of respiratory sample types (also including bronchoalveolar lavage, sputum and saliva), which may yield different degrees of contaminating bacterial and/or human reads. Loss of assay sensitivity due to the presence of high-level background DNA from either host or bacterial origin is a fundamental issue for metagenomic approaches, even in cell free sample types such as cerebrospinal-fluid [36]. This challenge is exacerbated in throat swabs, as seen in our data. Our use of hazara virus as an internal positive control allows us to identify those sample in which sensitivity has dropped below 10^4^ viral genome copies per ml. In our test set 8% of samples showed insufficient sensitivity for hazara virus, however half of these contained a high titre of influenza, so only 4% were true sensitivity failures. This figure is in line with the reported 6% failure rate due to high background for RNA virus detection from a clinically validated metagenomic sequencing assay for pathogen detection in cerebrospinal-fluid [36]

At the higher Ct values in our clinical samples (Ct 30-40), the sensitivity of Nanopore sequencing was reduced compared to the current PCR-based test (Cepheid). Further optimisation will be required to maximise the diagnostic yield from this group of samples, without sacrificing specificity.

The correlation between Ct value and Nanopore reads confirms semi-quantitative output. Using samples from the ferret influenza model, collected under standardized laboratory conditions, demonstrated excellent reproducibility of viral read proportions at a given viral titre across biological replicates. However, we observed heterogeneity in output between clinical samples as well as between Nanopore flowcells, suggesting that the current platform is not yet sufficiently reliable for reproducibly generating quantitative data. In addition, the detection of positive controls can be impaired in high background samples also creating further uncontrolled variability in the test yield.

Future application of this method will involve real-time laboratory testing of respiratory samples, running the platform head-to-head with existing clinical diagnostics to further assess sensitivity and specificity, and using influenza sequence data to investigate transmission events. Identifying instances of nosocomial transmission may shed light on healthcare-acquired infection, thus helping to improve infection control practice. Assessment of diversity within deep sequence datasets provides an opportunity to investigate the relationship between within-host polymorphisms and clinical outcomes. Long-read sequences confer the potential advantage of identifying viral haplotypes and ascertaining the extent to which significant polymorphisms are transmitted together or independently [25]. We have shown that the method is robust for the identification of commonly circulating influenza strains in human populations, but further investigation is required to ascertain the extent to which it performs reliably in other (avian, animal) strains.

In summary, our methods show promise for generating influenza virus sequences directly from respiratory samples. The ‘pathogen agnostic’ metagenomic sequencing approach offers an opportunity for simultaneous testing for a wide range of potential pathogens, providing a faster route to optimum treatment and contributing to antimicrobial stewardship. Longer term, this approach has promise as a routine laboratory test, providing data to inform treatment, vaccine design and deployment, infection control policies, and surveillance.

## Methods

### Study cohort and sample collection

We collected respiratory samples from the clinical microbiology laboratory at Oxford University Hospitals NHS Foundation Trust, a large tertiary referral teaching hospital in the South East of England. We worked with anonymised residual material from throat and nose swabs generated as a result of routine clinical investigations, between January and May 2018. Samples were collected using a sterile polyester swab inoculated into 1-3 ml of sterile viral transport medium (VTM), using a standard approach as described on the CDC website [37]. Workflow is shown in Fig 1.

During the study, respiratory samples submitted to the clinical diagnostic laboratory were routinely tested by a PCR-based test using GeneXpert (Cepheid) to detect influenza A and B, and respiratory syncytial virus (RSV). Samples from patients in designated high-risk locations (haematology, oncology and critical care) were tested using Biofire FilmArray (Biomérieux) to detect an expanded panel of bacterial and viral pathogens. Quantitative data (Cycle threshold, Ct) were generated by the GeneXpert, and we used the Influenza Ct value to estimate viral titre in clinical samples. Using GeneXpert, up to 40 PCR cycles are performed before a sample is called negative (ie positives have Ct <40). Quantification was not available for Biofire results.

For methodological assessment, we focused on four categories of samples:

- Positive pool: we pooled 19 throat swab samples that had tested positive for influenza A in the clinical diagnostic laboratory to provide a large enough sample to assess reproducibility (Fig 1B);
- Negative pools: we generated three pools of throat swab samples that had tested negative for influenza (consisting of 24, 38 and 38 individual samples respectively) (Fig 1B);
- Individual positive samples: we included 40 individual samples (35 throat swabs and 5 nasal swabs) that had tested positive for influenza A or B, selected to represent the widest range of GeneXpert Ct values (13.5 to 39.3; valid test result range 12 to 40).
- Individual negative samples: we selected 10 individual throat swab samples that were influenza negative.

### Quantification of viral RNA in samples

We quantified viral titres in Hazara virus stocks and pooled influenza A positive throat swabs by qRT-PCR using previously described assays and standards [38,39].

### Optimisation of methods

Prior to establishing the protocol detailed in full below, we assessed the impact of two possible optimisation steps, as follows:

i. Centrifugation vs filtration: We investigated two approaches to depleting human/bacterial nucleic acid from our samples, filtration of the raw sample via a 0.4μm filter (Sartorius) before further processing, versus a hard spin (16,000 xg for 2 min). cDNA libraries for this comparison were produced as described previously [21].
ii. Reduced time cDNA synthesis: To assess the possibility of time-saving in the cDNA synthesis steps, we compared performance of the previously described protocol [21] to a modified version with two alterations, first using SuperScript IV (ThermoFisher) in place of Superscript III (ThermoFisher) for reverse transcription with incubation time reduced from 60min to 10min at 42°C, and secondly reducing the cDNA amplification PCR extension cycling time from 5min to 2min.

### Positive control

Prior to nucleic acid extraction, each sample was spiked with Hazara virus virions to a final concentration of 10^4^ genome copies per ml as a positive internal control. This is an enveloped negative-stranded RNA virus (genus Orthonairovirus, order Bunyavirales) with a tri-segmented genome of 11980, 4575 and 1677 nucleotides in length (Genbank KP406723-KP406725). It is non-pathogenic in humans, and would therefore not be anticipated to arise in any of our clinical samples. Cultured virions from an SW13 cell line were provided by the National Collection of Pathogenic Viruses (NCPV), catalogue no. 0408084v).

### Nucleic acid extraction

Samples were centrifuged at 16,000 xg for 2min. Supernatant was eluted, without disturbing pelleted material, and used in nucleic acid extraction. Total nucleic acid was extracted from 100 μl of supernatant using the QIAamp viral RNA kit (Qiagen) eluting in 50ul of H_2_O, followed by a DNase treatment with TURBO DNase (Thermo Fisher Scientific) at 37·C for 30min. RNA was purified and concentrated to 6μl using the RNA Clean & Concentrator™-5 kit (Zymo Research) following manufacturer’s instructions. Randomly amplified cDNA was prepared for each sample using a Sequence Independent Single Primer (SISPA) approach, adapted from our previously described workflow [21], based on the Round A/B methodology [24]. For reverse transcription, 4μl of RNA and 1 μl of Primer A (5’-GTTTCCCACTGGAGGATA-N9-3’, 40 pmol/μl) [24] were mixed and incubated for 5min at 65°C, then cooled to room temperature. First strand synthesis was performed by the addition of 2μl SuperScriptIV First-strand Buffer, 1μL 12.5mM dNTPs, 0. 5μL 0.1 M DTT, 1μl H_2_O, and 0.5μL SuperScriptIV (ThermoFisher) before incubation for l0min at 42°C. Second strand synthesis was performed by addition of 1μl Sequenase Buffer, 3.85μl H_2_O and 0.15μl Sequenase (Affymetrix) prior to incubation for 8min at 37°C, followed by addition of 0.45μl Sequenase Dilution Buffer, 0.15μl Sequenase and a further incubation at 37°C for 8min. Amplification of cDNA was performed in triplicate using 5μl of the reaction as input to a 50μl AccuTaq LA reaction (Sigma) according to manufacturer’s instructions, using 1μl Primer B (5’-GTTTCCCACTGGAGGATA-3’) [24], with PCR cycling conditions of: 98°C for 30s; 30 cycles of 94°C for 15s, 50°C for 20s, and 68°C for 2min, followed by 68°C for 10min. Amplified cDNA was pooled from the triplicate reactions and purified using a 1:1 ratio of AMPure XP beads (Beckman Coulter, Brea, CA) and quantified by Qubit High Sensitivity dsDNA kit (Thermo Fisher), both according to manufacturer’s instructions.

### Nanopore Library Preparation and Sequencing

Multiplex sequencing libraries were prepared using 250ng of cDNA from up to 12 samples as input to the SQK-LSK108 or SQK-LSK109 Kit, barcoded individually using the EXP-NBD103 Native barcodes (Oxford Nanopore Technologies) and a modified “One-pot” protocol (Quick 2018). Libraries were sequenced on FLO-MIN106 flow cells on the MinION Mk1b or GridION device (Oxford Nanopore Technologies), with sequencing proceeding for 48h. Samples were batched according to the GeneXpert Ct value (see supplementary data file 1).

### Illumina methods

Nextera XT V2 kit (Illumina) sequencing libraries were prepared using 1.5ng of amplified cDNA as per manufacturer’s instructions and sequenced on a 2×150 bp-paired end Illumina MiSeq run, by Genomics Services Development Unit, Public Health England. Read mapping and data analysis as described previously [21]

### Bioinformatic analysis

Nanopore reads were basecalled using Guppy (Oxford Nanopore Technology, Oxford, UK). Output basecalled fastq files were demultiplexed using Porechop v0.2.3 (https://github.com/rrwick/Porechop). Reads were first taxonomically classified against RefSeq database using Centrifuge v1.0.3. Reads were then mapped against the reference sequence selected from Centrifuge report using Minimap2 v2.9. A draft consensus sequence was generated by using a majority voting approach to determine the nucleotide at each position. The resulting draft consensus sequences were BLASTed against a flu sequence database that included >2,000 H1N1 and H3N2 seasonal influenza sequences between 2018 and 2019, and downloaded from the Influenza Research Database [40]. Reads were again mapped against the reference sequence using Minimap2 v2.9 and the number of mapped reads were calculated using samtools v1.5 and Pysam (https://github.com/pysam-developers/pysam). The subtype of the influenza A virus derived from each clinical sample was determined by the subtype of HA and NA reference sequence. Consensus sequence was built using Nanopolish v0.11.0 and margin_cons.py script (https://github.com/zibraproject/zika-pipeline).

### Ferret study

We applied our sequencing approach to residual samples collected in a previous time-course experiment undertaken in a controlled laboratory environment [30]. We tested ferret nasal saline wash samples from three independent animals over an eight day time course, from three days prior to first exposure with influenza H1N1pdm09 and at days 1, 2, 3 and 5 post-infection. Sampling and plaque assays of viral titre were described previously [30].

### Ethics approval

The study of anonymised discarded clinical samples was approved by the London - Queen Square Research Ethics Committee (17/LO/1420). Ferret samples were residual samples from an existing study [30], for which the project license was reviewed by the local Animal Welfare and Ethics Review Board of Public Health England (Porton) and was subsequently granted by the Home Office.

### Data availability

Following removal of human reads, our sequence data have been uploaded to the European Bioinformatics Institute https://www.ebi.ac.uk/, project reference PRJEB32861.

### Funding

The study was funded by the NIHR Oxford Biomedical Research Centre. Computation used the Oxford Biomedical Research Computing (BMRC) facility, a joint development between the Wellcome Centre for Human Genetics and the Big Data Institute supported by Health Data Research UK and the NIHR Oxford Biomedical Research Centre. The views expressed in this publication are those of the authors and not necessarily those of the NHS, the National Institute for Health Research, the Department of Health or Public Health England, PCM is funded by the Wellcome Trust (grant ref 110110). DWC, TEAP and ASW are NIHR Senior Investigators.

## Supplementary material

**Figure S1.**
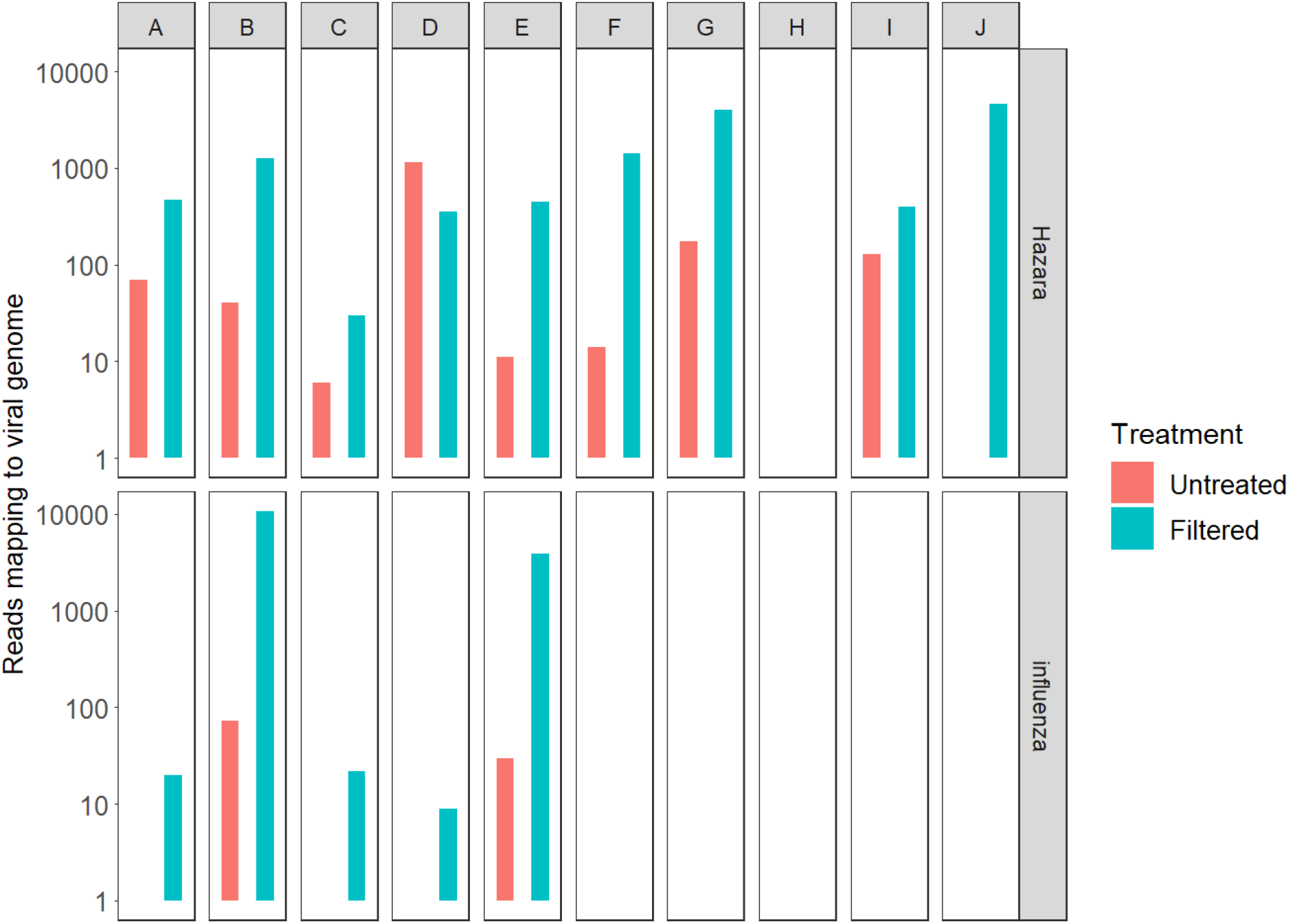
Comparison of viral read detection in throat swab samples with and without a filtration step. Five influenza A positive throat swab samples (A-E) and five negative samples (F-J) were spiked with Hazara virus as positive control at 10^4^ genome copies/ml and sequenced with and without filtration via a 0.4μm filter prior to RNA extraction. Reads mapping to influenza or Hazara are shown for each sample.

**Figure S2.**
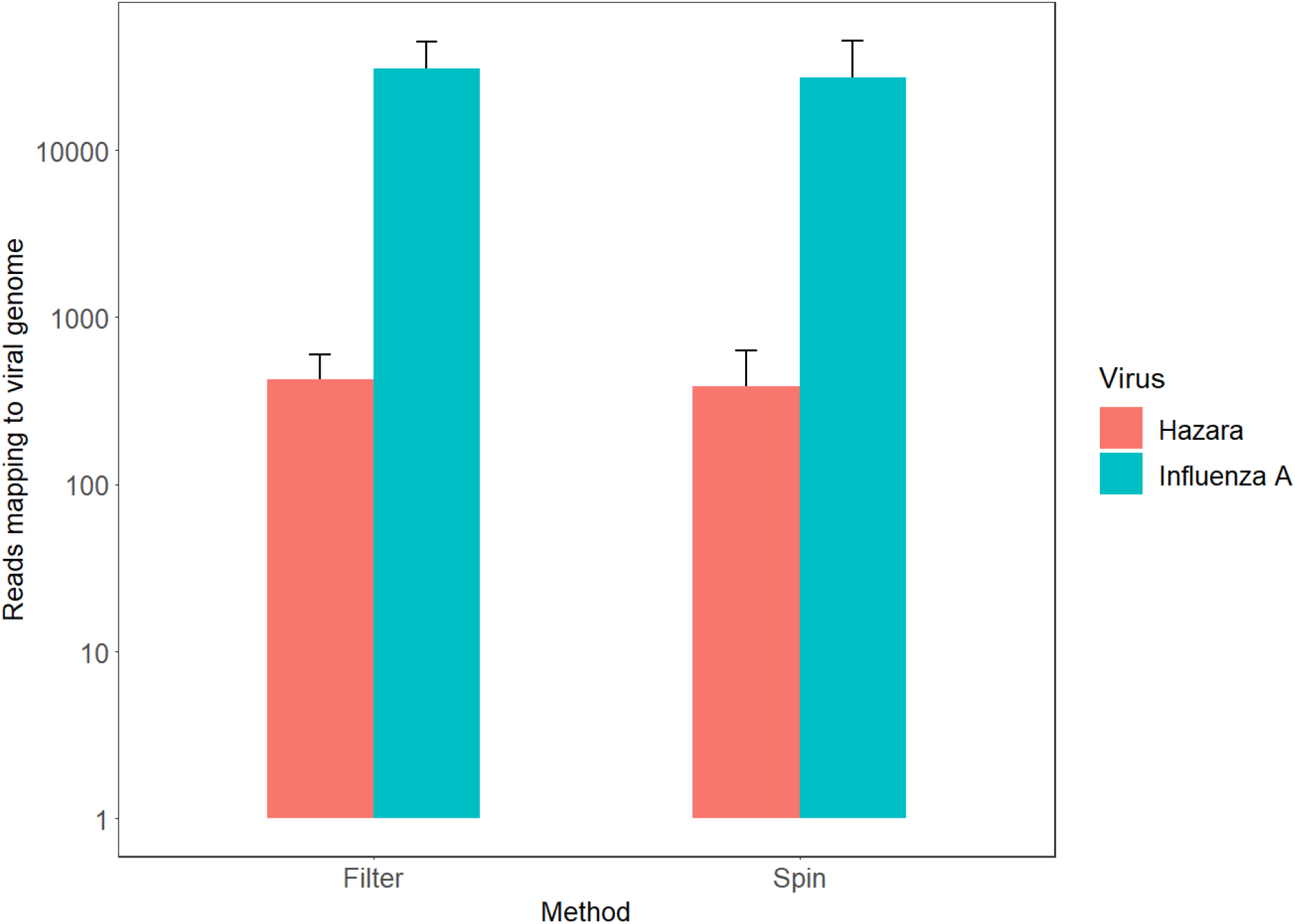
Comparison of Filtration and Centrifugation as Pre-extraction treatments to improve viral detection. Using a pool of 19 influenza A positive samples as input, triplicate extractions utilising either supernatant separation by centrifugation at 16,000 xg for 2 min, or filtration via a 0.4μm filter, were processed and all samples sequenced on a single flow cell. Mean total read numbers for each virus are plotted for each treatment, error bars indicate standard deviation.

**Figure S3.**
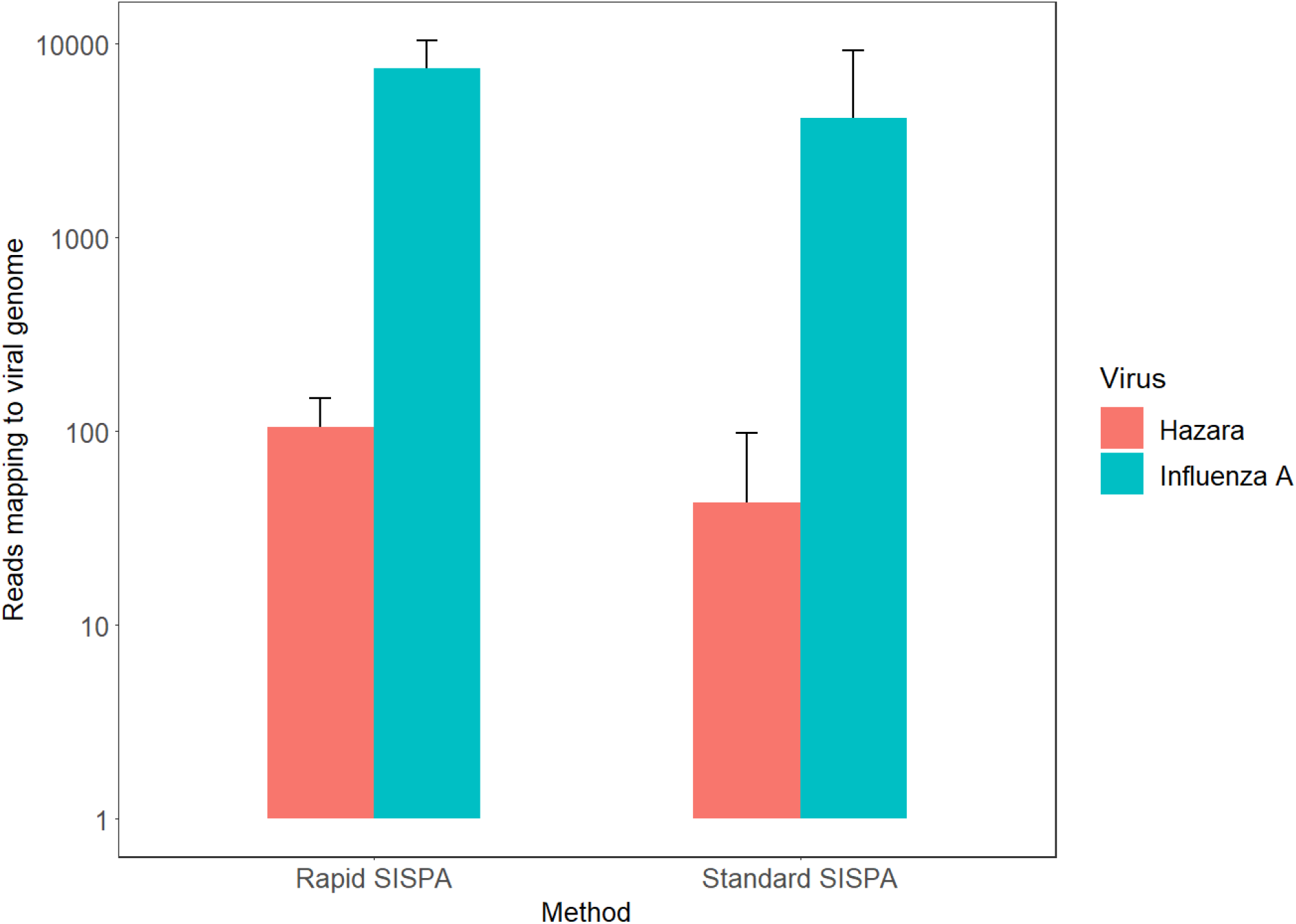
Assessment of effect of original and reduced processing time method on metagenomic viral sequencing. Using a pool of influenza A positive samples as input, triplicate extractions were processed by either existing (standard) or reduced incubation times (rapid) method and all samples sequenced on a single flow cell. Mean total read numbers for each virus are plotted for each treatment, error bars indicate standard deviation.

**Figure S4.**
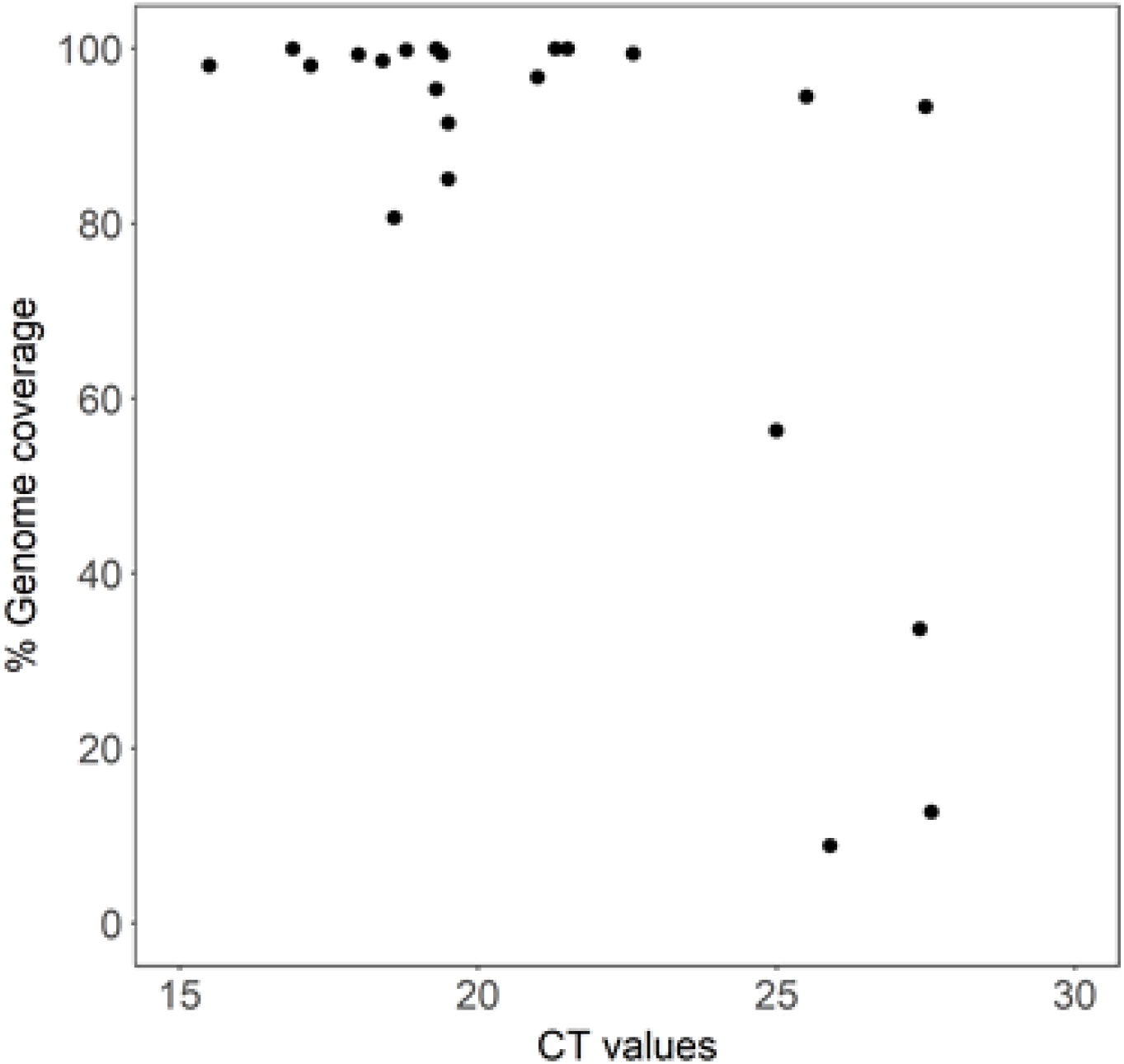
Coverage of influenza sequences recovered by nanopore metagenomic sequencing. Ct values were derived by testing using GeneXpert (Cepheid) in a clinical diagnostic laboratory. Genome coverage shown is the proportion of bases of the reference sequence that were called in the final consensus sequence for each sample.

**Figure S5.**
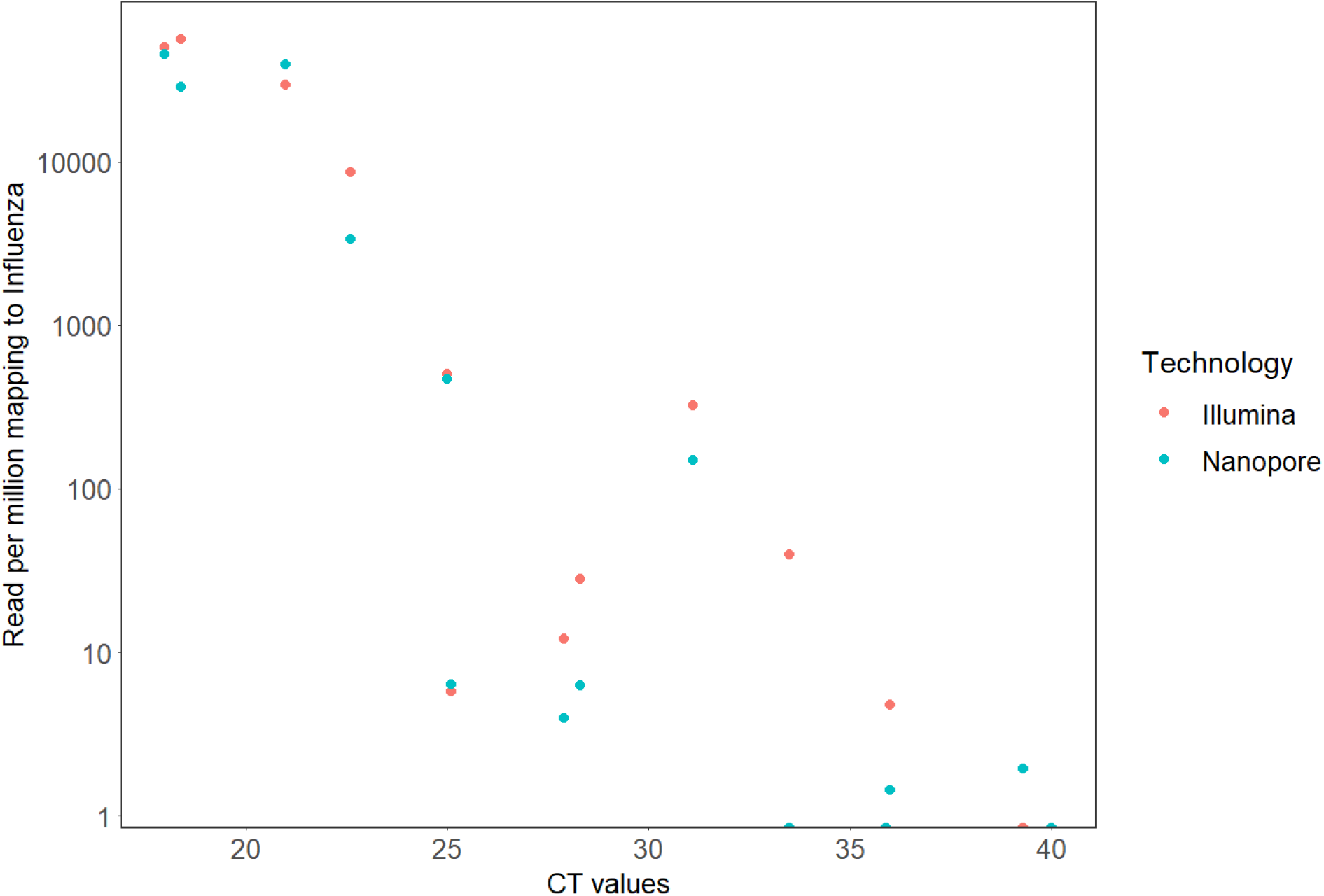
Comparison of proportions of influenza reads derived by Nanopore or Illumina sequencing for a subset of the individual samples across a range of Ct values. Ct values were derived by testing using GeneXpert (Cepheid) in a clinical diagnostic laboratory. Proportions are shown as number of influenza reads per million reads.

**Table S1. Summary data for the 50 individual throat swab samples.**

